# Additive Manufacturing of real-scale carotid artery models: The content–container interaction in sickle cell disease-related cerebral vasculopathy

**DOI:** 10.1101/2025.11.09.687432

**Authors:** Saskia Eckert, Christian Kassasseya, Morgane Garreau, Curie Jordane Kakpovi, Irène Vignon-Clementel, Suzanne Verlhac, Blanche Bapst, Luca Scarcia, Henri Guillet, Katy Dremont, Smaïne Kouidri, Frédéric Segonds, Kim-Anh Nguyen- Peyre, Pablo Bartolucci

## Abstract

Sickle cell disease–related cerebral vasculopathy depends on patient-specific vascular geometry and hemodynamics that are not captured by conventional experimental systems or animal models. To address this limitation, we developed an additively manufactured artery scaffold representing the container and integrated it into a controlled fluidic circuit reproducing physiological flow profiles, pressures, and viscosities, the content. Validation with clinical imaging confirmed the anatomical accuracy and flow fidelity of the additively manufactured models. Mechanosensitive cells, including endothelial cells and platelets, were incorporated into the model for biological analysis. This study serves as a proof of concept demonstrating how the Container–Content model can enhance the mechanistic understanding of vascular biology under physiologically relevant conditions. The ethically responsible platform bridges computational simulations and in vitro experimentation, offering a versatile foundation for investigating cerebrovascular complications in sickle cell disease and advancing the field of personalized medicine.

## Introduction

Sickle cell disease (SCD) is the most common severe monogenic disorder worldwide [1], with approximately 300,000 newborns affected each year [2]. It results from a β-globin gene mutation producing hemoglobin S [3], which polymerizes when deoxygenated, deforming RBCs leading to hemolysis, damaging vessels, and causing multi-organ injury.

SCD-related cerebral vasculopathy (CV), typically arising in childhood, is the first cause of pediatric stroke, including overt and silent strokes leading to cognitive impairment, disability, or death [4]. Stroke risk correlates with anemia and elevated cerebral blood flow (CBF) velocities >200 cm/s, measured by transcranial Doppler (TCD) ultrasound [4], [5], [6]. Pathogenesis involves hemorheological, biological, and genetic susceptibility mechanisms, notably endothelial inflammation and subsequent dysfunction of vascular cells [7],[8], [9], [10]. Stenoses of large arteries, particularly in the internal carotid artery (ICA) siphon, middle cerebral artery (MCA), and anterior cerebral artery (ACA), are commonly visualized by MRI, motivating our focus on the ICA segment for targeted modeling and analysis.

SCD-related CV thus represents a showcase pathology of content and container, in which altered blood rheology, with circulating elements such as hemolysis products and platelet-derived factors (content), interacts with a dysfunctional vascular wall (container), driving progressive endothelial injury and vascular remodeling.

Our previous works demonstrated that intracranial velocity accelerations preceding arterial lesions cannot be explained by a single anatomical factor but by the combined effect of flow rate and vascular geometry, including diameter, curvature, etc.., quantified by the Dean number [11]. Such accelerations could generate disturbed flow conditions in the CA, which in some vascular pathologies, may promote stenosis resulting from endothelial injury and intimal proliferation of smooth muscle cells (SMC) [12]. These disturbed flow patterns illustrate how mechanical imbalances within the content translate into structural alterations of the container.

To replicate these complex conditions, bioreactors and organ-on-chip systems enable precise flow control in anatomically relevant vessels, linking personalized hemodynamics with cellular responses [13]. Animal models, particularly Berkeley and Townes mice [14], as well as porcine and ovine systems [15],[16], have significantly advanced SCD research. However, inter-species differences in blood flow rates and vascular geometry limit their translational accuracy [17]. The content (blood properties) and the container (vessel geometry) differ substantially across species, limiting direct extrapolation to human conditions. Nevertheless, existing vascular models remain limited in their ability to capture patient-specific geometries and clinically relevant hemodynamics. Organ-on-chip systems often employ simplified microchannels, while animal models fail to reproduce human CV. Thus, there is a pressing need for an in vitro approach that bridges clinical imaging and functional vascular biology.

To address this, Catano et al. developed a miniaturized patient-specific additively manufactured (aM) carotid model, including endothelial cells (ECs) and SMCs under constant flow, with hemodynamic similarity preserved via Buckingham-π scaling [18]. Additive Manufacturing (AM) refers to a set of techniques that build three-dimensional objects layer by layer, allowing precise fabrication of complex geometries tailored to individual patient anatomy.

Here, we present what is, to our knowledge, the first demonstration of a full-scale, patient-specific CA model, fabricated through AM of anatomically complex geometries, integrating pulsatile inflow and cellular components to enable spatially and temporally resolved hemodynamics at the blood–vessel wall interface. Unlike miniaturized organ-on-chip systems, our platform reproduces exact native vessel dimensions while integrating all relevant flow conditions, thereby providing a physiologically realistic environment for studying content–container interactions.

In this context, our approach directly couples the container and the content, allowing controlled investigation of their interface, which we define here as the EC layer. This layer is sensitive to mechanical forces, which are sensed via mechanoreceptors such as ion channels (notably Piezo1), G protein–coupled receptors, or membrane-associated structures [19], and also responds to soluble mediators released during platelet degranulation, hemolysis, and other stress-related processes.

Within the same framework, platelets represent a second player whose behavior is strongly regulated by hemodynamic conditions. They are mechanosensitive, activated by peak shear stress [20], exposure time [20], and shear gradient changes [21]. Repeated alternating shear increases sensitivity [22]. Platelet activation is typically accompanied by degranulation, releasing mediators such as TGF-β, thrombospondin (TSP), and serotonin, factors involved in vascular remodeling, inflammation, and endothelial dysfunction. These molecules are released in response to shear activation and interact with endothelial and SMC, thereby promoting vascular remodeling.

The goal of this study is to present an interdisciplinary and ethically responsible methodology to investigate how controlled blood flow and hemodynamic conditions affect ECs and platelets — the mechanosensitive cell types interacting at the arterial wall–blood interface (patent: WO2021/089665A1 [23]).

Our method integrates three main components:

1/ **The container:** A personalized artery model fabricated through AM;
2/ **The content:** From a hemodynamic perspective, it is defined as the fluidic system that reproduces controlled flow conditions, while from a biological perspective, it comprises mediators interacting with the interface, such as mechanosensitive circulating cells (platelets); and
3/ **The interface:** ECs acting as the communicator between the arterial wall and the blood flow (Figure 1A).

**Figure 1:**
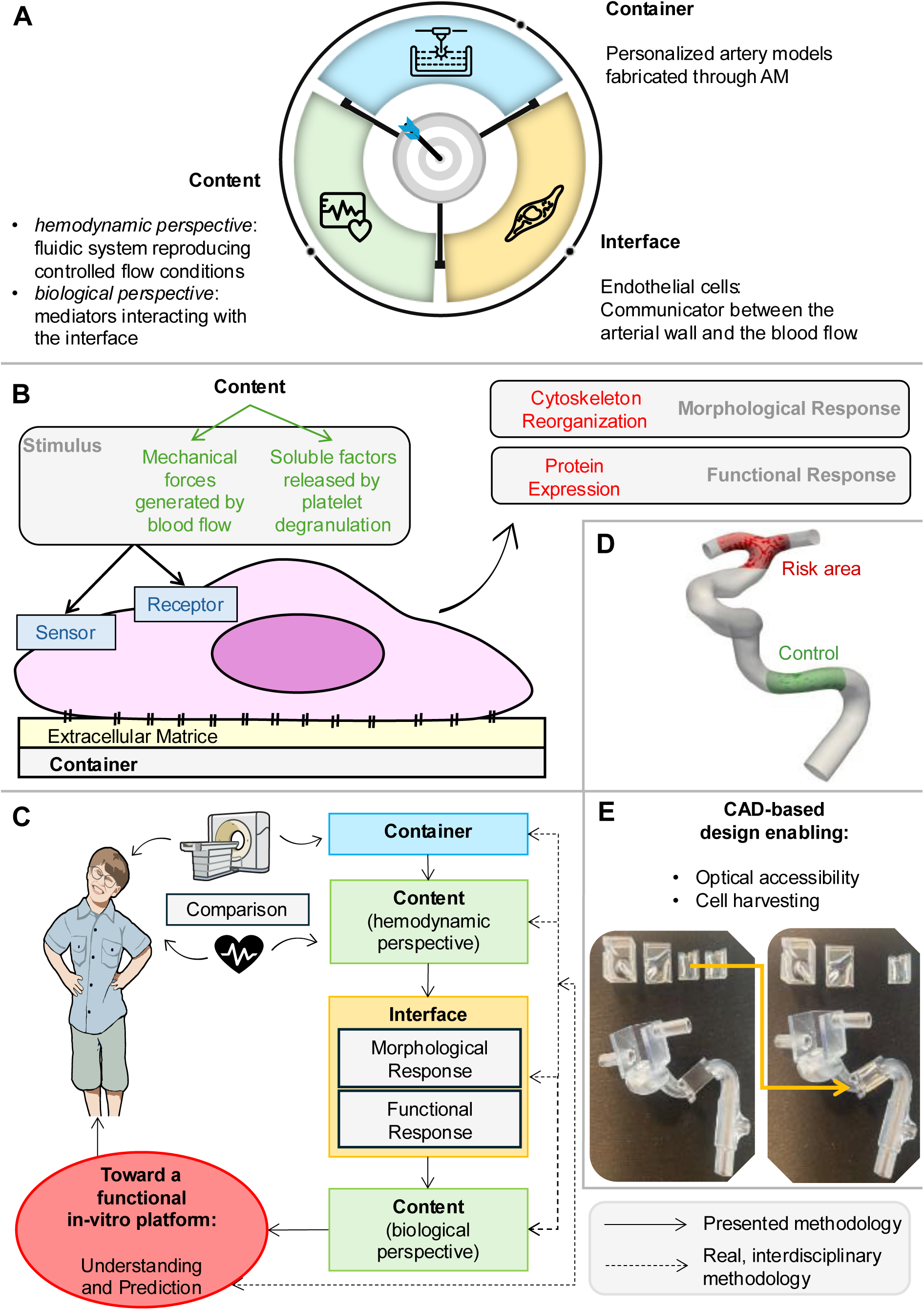
Schematic of the developed methodology (icons adapted from NIAID Visual & Medical Arts). A) Conceptual interplay between the Container, Content, and Interface model. B) Example of stimulus and corresponding endothelial cell responses at the Interface. C) Implementation of the interdisciplinary workflow towards a functional platform for vascular research. D) Risk and control zones in SCD carotid artery. E) Detachable and openable artery segments designed by Computer-Aided-Design (CAD).

Our AM-based approach integrates biological components with controlled hemodynamics to overcome the limitations of animal and conventional in vitro models, enabling precise investigation of vascular function under physiologically relevant conditions. By experimentally reconnecting the content and the container, this methodology establishes a physiologically meaningful bridge between hemodynamics and vascular biology in SCD-related vasculopathy.

## Results

The platform integrates mechanical and biological stimuli, linking them to EC morphological and functional responses under flow conditions and during vascular function or dysfunction (Figure 1B). An interdisciplinary workflow has been implemented to establish a functional system for vascular research (Figures 1C).

We present an female adult SCD case without known vasculopathy (39-year-old, 170 cm), using a flow rate of 480 mL/min [24]. The model targets the ICA, MCA, and ACA segments as a proof of concept for future applications. Viscosity is set to 4 cP, a value typically observed in SCD patients, which also maximizes sensitivity to hemodynamic effects and flow disturbances [25].

### Development of a real-scale carotid model integrating controlled hemodynamics: (The Content-Container model)

#### Additive Manufacturing of the carotid artery

AM, enabling fabrication from digitally designed models via Computer-Aided Design (CAD), provides design flexibility and precision in reproducing complex, patient-specific vascular geometries. We reconstruct the vessel lumen and outer wall separately, with the outer wall modeled in parametric CAD software to ensure both design adaptability and preservation of patient-specific anatomy [26]. This digital approach enables targeted model customization, such as defining risk-prone regions (e.g., the carotid bifurcation, Figure 1D) or designing declipsable segments for spatially resolved cellular analysis (Figure 1E). Moreover, CAD facilitates the integration of functional features, such as connectors for flow control or data acquisition, and allows the creation of openable, flow-through architectures for direct visualization of endothelial behavior. This level of geometric control establishes CAD-based reconstruction as a transformative tool in the development of physiologically relevant, patient-specific vascular models.

#### Geometric validation of the aM carotid artery

The aM carotid model was scanned with an AngioScanner and compared to the patient’s DICOM image. The artery was suspended free in a box to avoid contact (Figure 3A). Additionally, a rotational 3D-DSA of the printed carotid model was acquired on a Siemens Artis zee fluoroscopy unit (Siemens Medical Solutions; Erlangen, Germany) (Figure 3B), and the resulting 3D reconstruction was directly compared with the original patient-derived 3D dataset.

Lumen curvature, bifurcation angulation and the carotid siphon morphology were preserved, confirming that the additively manufactured carotid reproduces the native patient anatomy with high geometric fidelity.

#### Integration of hemodynamic parameters into the AM carotid artery model: MCL

Pulsatile flow is reproduced with a Harvard Apparatus Pulsatile Blood Pump (Rabbit model, volume flow control, 65% diastolic duration) (Figure 2). The pump is placed at test-section height to avoid gravity-induced pressure drops. To reconstruct a typical flow of an adult sickle cell patient (480 mL/min), the heart rate was set to 80 bpm and the stroke volume to 6 mL. Low-compliance tubing connects the pump to a compliance chamber, then to the carotid entry. Downstream resistance is set by a needle-pinch valve at the MCA/ACA Y-piece (equal flow split assumed). Needle-pinch valves provide precise control and sterile handling.

**Figure 2:**
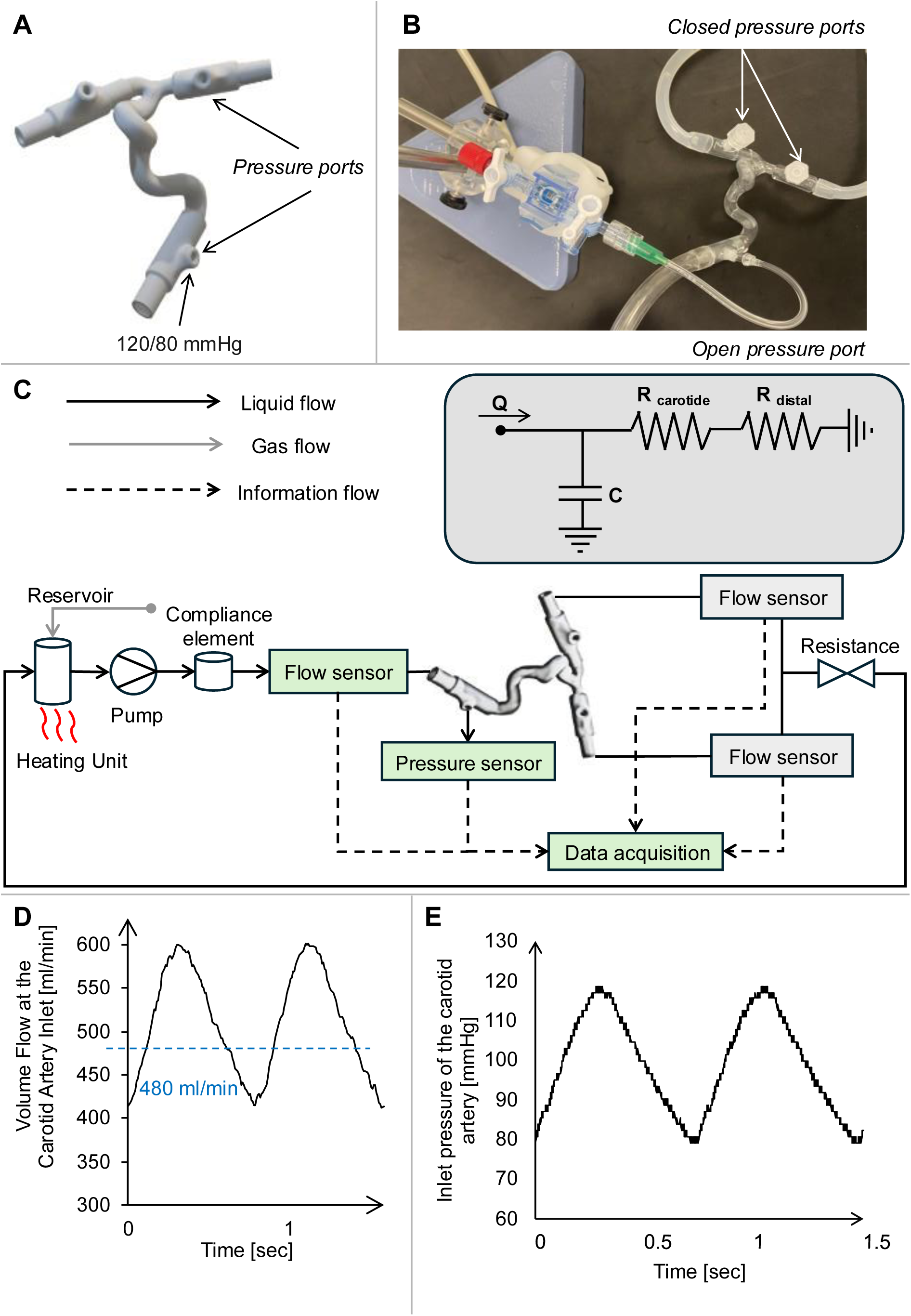
Peripheral setup for reconstructing personalized flow in the aM carotid artery. A) Printed artery with integrated pressure ports. B) Open and sealed ports (closable after flow adjustment). C) Fluidic circuit and electrical analogue. D) Flow waveform at 80 bpm (stroke volume 6 ml, total flow 480 ml/min, viscosity 4 cP). E) Corresponding pressure profile.

**Figure 3:**
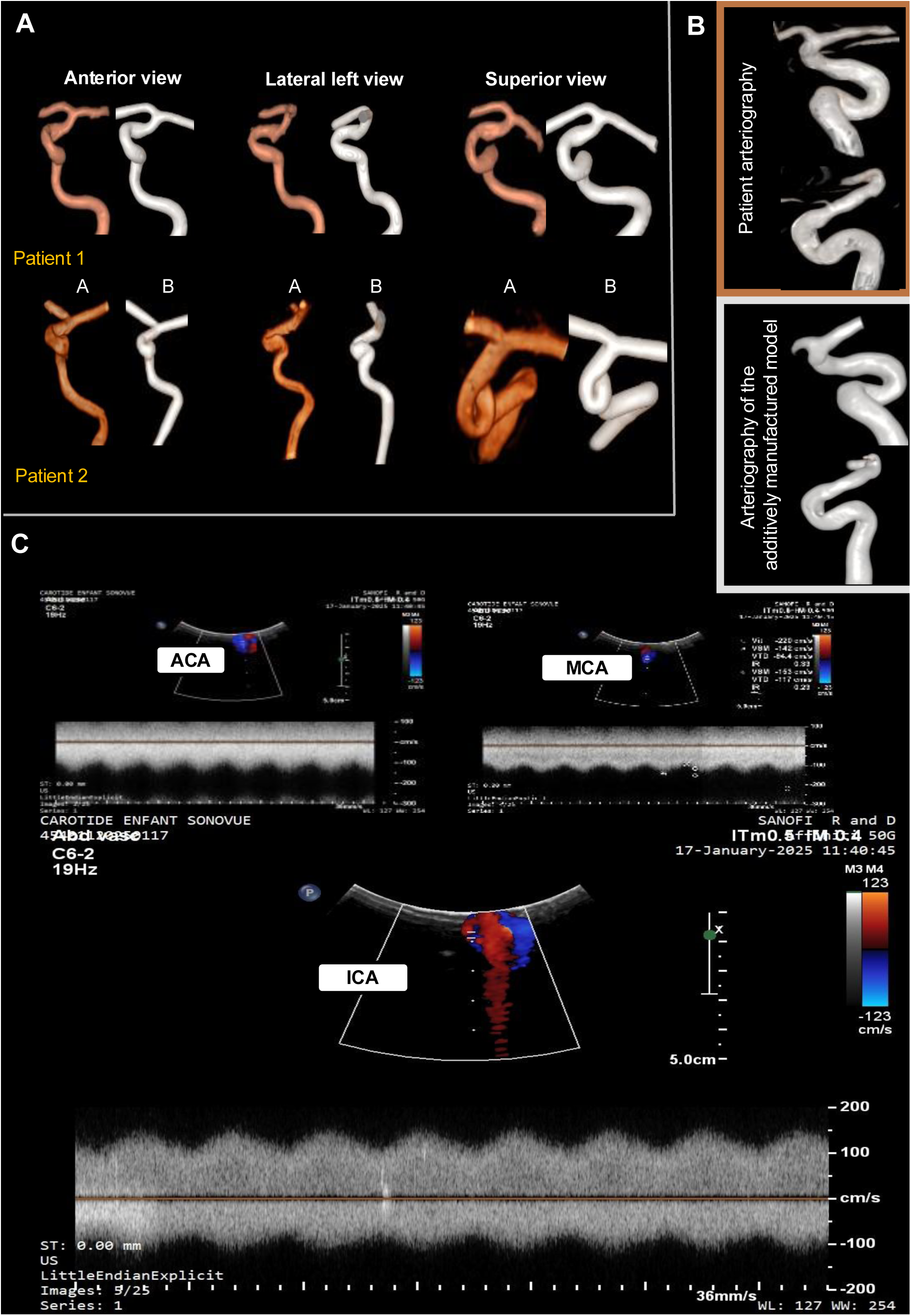
Validation with existing imaging and diagnostic modalities. A) 3D volume-rendered views from in vivo ToF-MR and corresponding post-contrast CT acquisitions of aM carotids. B) 3D arteriography volume rendering from the patient compared to the aM model. C) Doppler ultrasound measurements in the ICA, MCA, and ACA segments.

Dextran MW 150,000 (Sigma-Aldrich) adjusts viscosity to 4cP. Complete EBM-2 medium (Lonza) is first mixed with 17.5% dextran, sterile-filtered (0.2 µm), then diluted to 5.1% dextran. This minimizes filtration of the final medium. The mixture is vortexed and left overnight until fully dissolved. At high shear, dextran medium behave as Newtonian fluids.

Volume flow is measured with an FD-XC8R1 flow meter (Keyence, Japan) connected via a DFROBOT current–voltage converter to an Arduino Uno R3. Pressure ports are directly printed in the carotid entry. Pressures are recorded with an APT300 transducer (Harvard Apparatus) linked to a TAM-A PLUGSYS amplifier (Hugo Sachs, Germany) and monitored with a SmartScope oscilloscope (LabNation). The results demonstrated a pressure drop of 0.6 mmHg per mm of tubing between the transducer and the artery entry was measured and corrected. Direct CO₂ injection into the flask supplying the pump with dextran-adapted medium was controlled at 5%, enabling gas exchange at the liquid surface (Figure 2). Medium is maintained at 37°C.

#### Fluidic and hemodynamic validation

A color Doppler ultrasound machine equipped with a 6-2 MHz convex probe was used for validation. After viewing the geometry of the model using color Doppler imaging, flow velocity waveforms of the MCA, ACA and ICA were obtained using pulsed Doppler with an insonation angle of 0°. TUSK XC2700T material is acoustically transparent. Water viscosity was set to 4 cP with Dextran 150,000 MW. SonoVue microbubbles (SF₆) served as diagnostic contrast, enhancing signal reflection (Figure 3C).

In our model, Time-Average Mean Maximum velocity (TAMMV) in the ICA reaches 125 cm/s and systolic peak 150 cm/s. For pediatric patients, a TAMM velocity of 170 cm/s typically indicates the need for short-term follow-up [27]. Thus, our model reproduces a low-resistance pulsatile flow comparable to the patient’s flow with clinically relevant velocities without indicating a transcranial Doppler–based risk of stroke. However diastolic velocity is slightly higher than typically observed in vivo (∼80 cm/s) with a lower systolic-diastolic ratio and a smooth sinusoidal waveform in contrast to physiological waveforms, which feature a sharper systolic upstroke and more pronounced diastolic deceleration due to native vessel compliance and resistance. This behavior arises from the simplified two-element Windkessel integrated into our Mock Circulatory Loop (MCL), consisting of a pump and downstream compliance and resistance elements [28],[29]. While more complex three-or four-element Windkessel models can better capture flow impedance and vascular inertia [30],[31], we prioritized simplicity to facilitate experimental handling and reproducibility.

### Endothelialization of the aM carotid artery within the MCL (container–interface)

#### Endothelial Cell Seeding

ECs line blood vessels in many organs and form the interface between arterial wall and blood flow, exhibiting structural and functional heterogeneity [32]. Arterial ECs are elongated, have tight junctions, and adapt to high shear stress, showing minimal phenotypic change in response to inflammatory stimuli, whereas venous ECs are wider, more permeable, and strongly respond to inflammation [33],[34],[35]. It is hypothesized that the arterial EC phenotype varies along the arterial tree in response to local hemodynamics rather than its anatomical origin, as confirmed by single-cell RNA sequencing [36]. High shear stress can align venous ECs in vitro, demonstrating mechanosensitivity.

All parts are plasma-treated to obtain the hydrophilic surface then washed and sterilized. The detachable parts are coated with 0.5% gelatin and washed twice with PBS without Ca2+ et Mg2+. HUVECs (≤ passage 8) suspended in EBM-2 at 10⁶ cells/ml are seeded into the model. ECs from suspension attach to an extracellular matrix and can be guided using physical forces such as gravitational [37],[38],[39], magnetic [40],[41], hydrostatic [42], or combined forces [43]. ECs are seeded by gravity for 2 h, repeated three times with fresh cells and manual rotation for full coverage, then transferred to 6-well plates for 24 h proliferation (Figure 4A).

**Figure 4:**
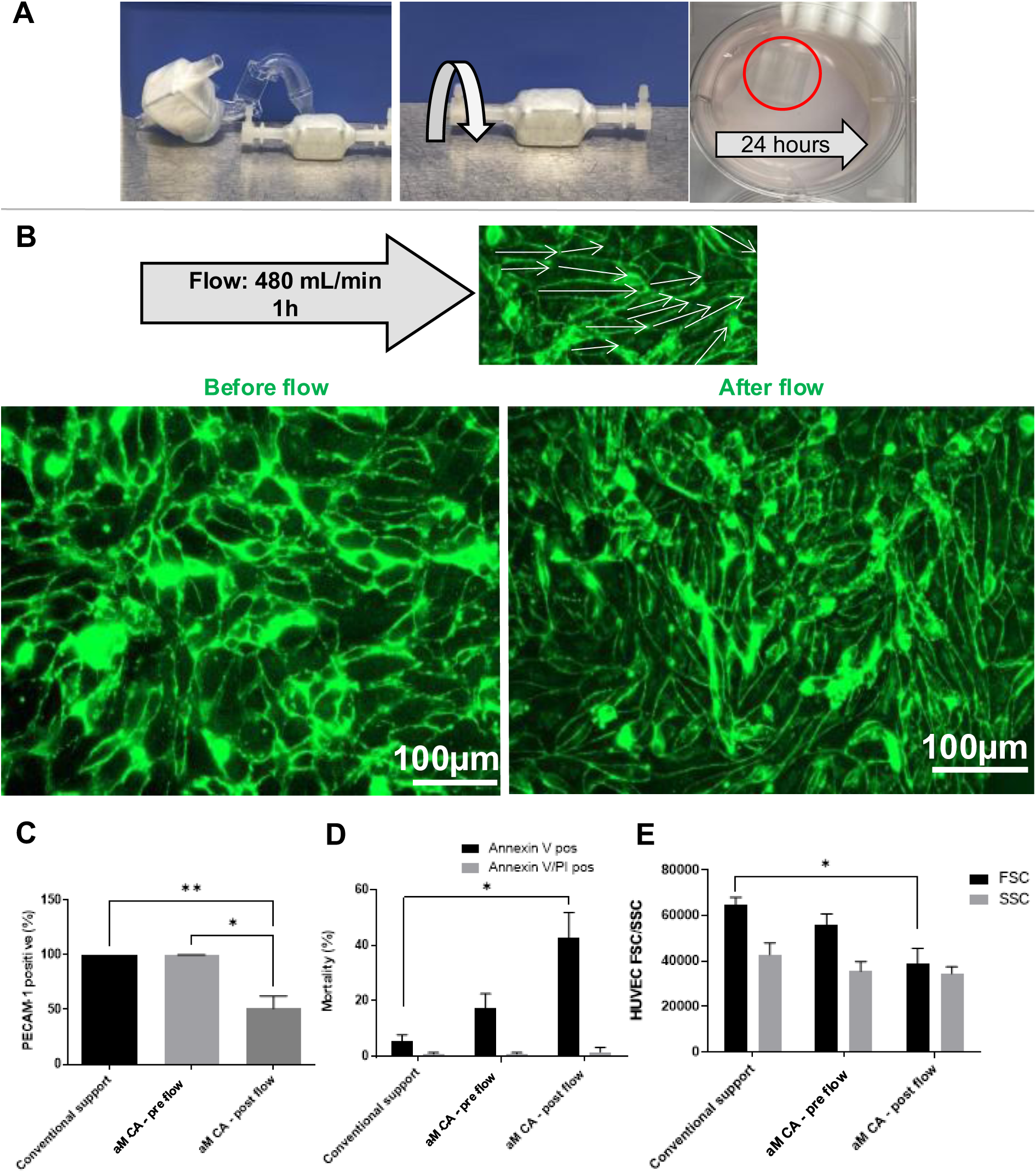
Integration of endothelial cells as the interface in the aM carotid model. A) Workflow: detachment, seeding, 24 h incubation. B) Comparison pre-/post-flow (Phalloidin, Z-stack projection) to assess morphological responses to flow conditions. C) Percentage of PECAM-positive and viable cells pre-/post-flow. D) Mortality pre-/post-flow with conventional support. E) FSC vs. SSC profiles pre-/post-flow with conventional support.

#### Integration of the detachable parts into the CA and insertion into the MCL

The endothelialized parts are reinserted into the aM shell model, which is then perfused for 1 h with the previously described MCL with 480 mL/min and a viscosity of 4cP. No pre-conditioning in pressure or gradual shear adaptation was applied.

#### Morphology of HUVEC in the aM CA

Cured TUSK (refractive index of 1.514) impairs phase-contrast imaging. Visualization of anti-PECAM-1 staining failed at various concentrations and wavelengths due to the high refractive index. Thus, highly fluorescent dyes are used.

ECs were fixed with 4% paraformaldehyde, permeabilized with Triton X-100, and stained with Hoechst to visualize nuclei and Phalloidin to label actin filaments. Because of the arterial curvature, Z-stacks were acquired in 3-µm steps using the ZEN microscope software. The high refractive index of the cured TUSK introduced background noise, which was minimized through image processing with Fiji or ZEN using a rolling-ball algorithm. The resulting stacks were combined by maximum intensity projection to confirm the presence and distribution of ECs, although the actin signal was less pronounced.

Cells were not uniformly aligned but showed preferential orientation. Given complex flow, comparison with simulated shear maps is needed. As vessel geometry was projected to 2D, no quantification (e.g., alignment factors) was done to avoid distortions affecting accuracy. Notably, alignment occurred after only 1h, whereas literature typically reports 24–48 h.

#### Inflammatory status and viability

Inflammatory status was assessed, enabling spatial visualization of activated cells in the CA model. ECs were fixed, stained with anti-ICAM-1 (activation), Hoechst (nuclei) then permeabilized for phalloidin staining (actin). Images pre-/post-flow showed TNFα increased ICAM-1 expression on HUVECs, confirming direct visualization of endothelial activation for future spatial analyses (Figure 5A).

**Figure 5:**
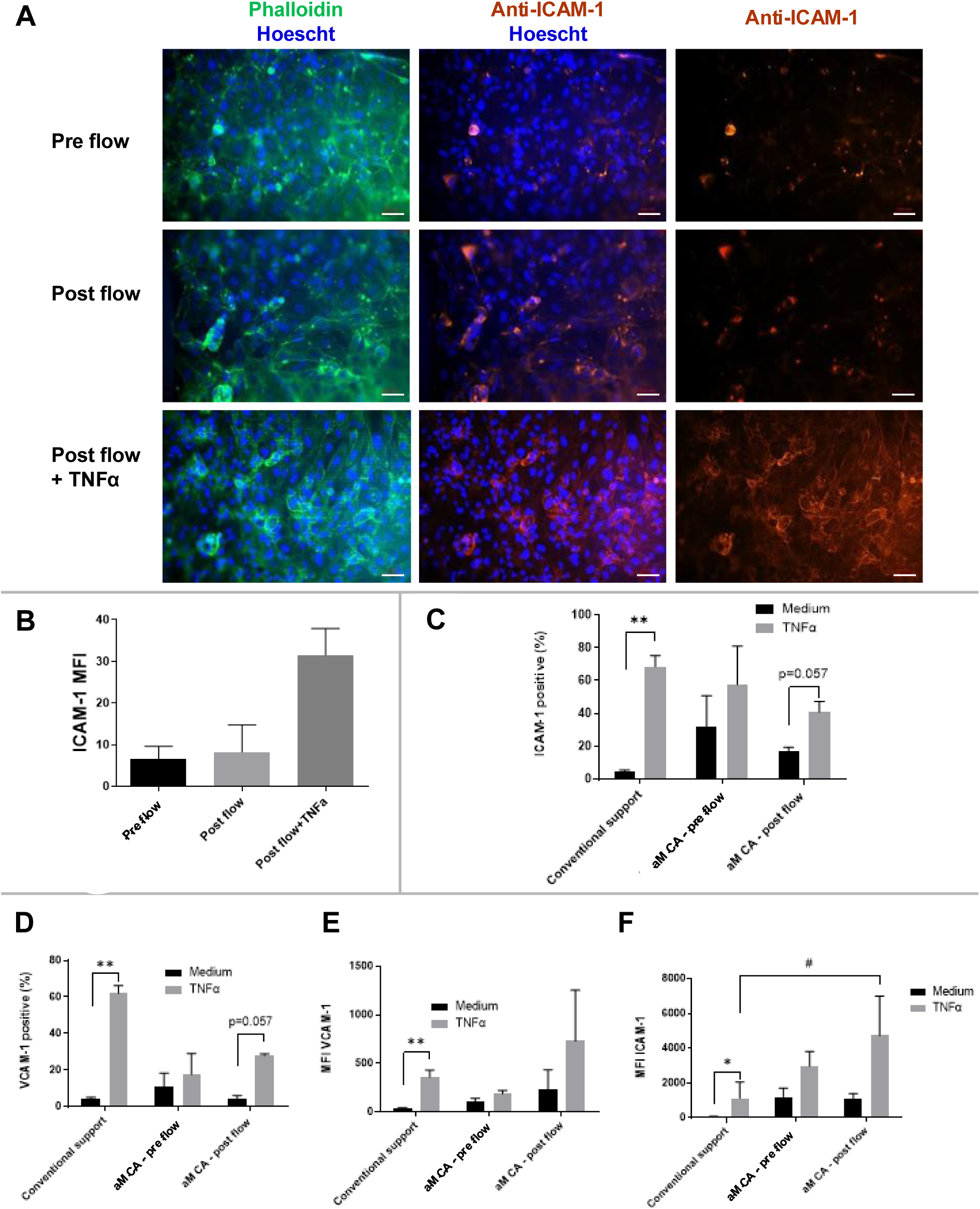
Inflammatory responses of the endothelial cell interface. A) Microscopic analysis of ICAM-1–positive cells for spatial localization of inflammation. B) Mean fluorescence intensity (MFI), n = 3 experiments, 5 images per experiment. C) ICAM-1– positive cells detected by flow cytometry. D) VCAM-1–positive cells detected by flow cytometry. E) MFI of VCAM-1 from flow cytometry. F) MFI of ICAM-1 from flow cytometry.

Detachable CA parts pre-and post-flow exposure were compared with conventional cultures on multiwell plates. CA parts required higher seeding density to reach confluence within 24h. HUVECs were subsequently trypsinized, washed and stained for flow cytometry.

PECAM-1, an endothelial marker, was significantly decreased by 50% after flow exposure suggesting that direct arterial pulsatile flow without preconditioning may cause endothelial damage or PECAM-1 shedding (Figure 4C).

Cell viability was assessed by Annexin V/PI staining (Figure 4D). The percentage of Annexin V–positive cells increased following flow exposure (42.6% vs 17.4%), while the percentage of Annexin V/PI double-positives were unchanged, indicating early apoptosis. FSC decreased and SSC remained stable, consistent with an early apoptotic phenotype. Such direct arterial pulsatile flow (480 mL/min) without preconditioning thus induces early apoptosis without widespread cell death (Figure 4E).

Flow cytometry analysis of ICAM-1 and VCAM-1 expression revealed no differences between pre-and post-flow conditions, indicating that physiological flow alone does not activate ECs within the model. As a positive control for endothelial activation, TNF-α stimulation markedly increased the proportion of ICAM-1–positive cells in both pre-and post-flow conditions, with a higher mean fluorescence intensity (MFI) after flow exposure (Figure 5B). These results suggest that ECs remain responsive to TNF-α stimulation after flow, and that prior flow exposure may enhance the magnitude of activation without altering the fraction of responsive cells (Figure 4C-F).

#### Platelet Apheresis Concentrate experiments in aM carotid model (Content)

The same CA model was perfused with 200 mL platelet apheresis concentrate (APC) at physiological (480 mL/min) or high flow rates (800 mL/min) to study platelets activation and degranulation over time, with samples collected at 0, 5, 15, and 30 min (n = 3, both flow conditions used the same donor). Platelets experiments were performed at room temperature (22°C).

Platelet counts in APC were performed pre-and post-perfusion to monitor losses. Fold change showed no significant change under either flow condition, indicating negligible loss and no adhesion to the vessel wall (Figure 6A).

**Figure 6:**
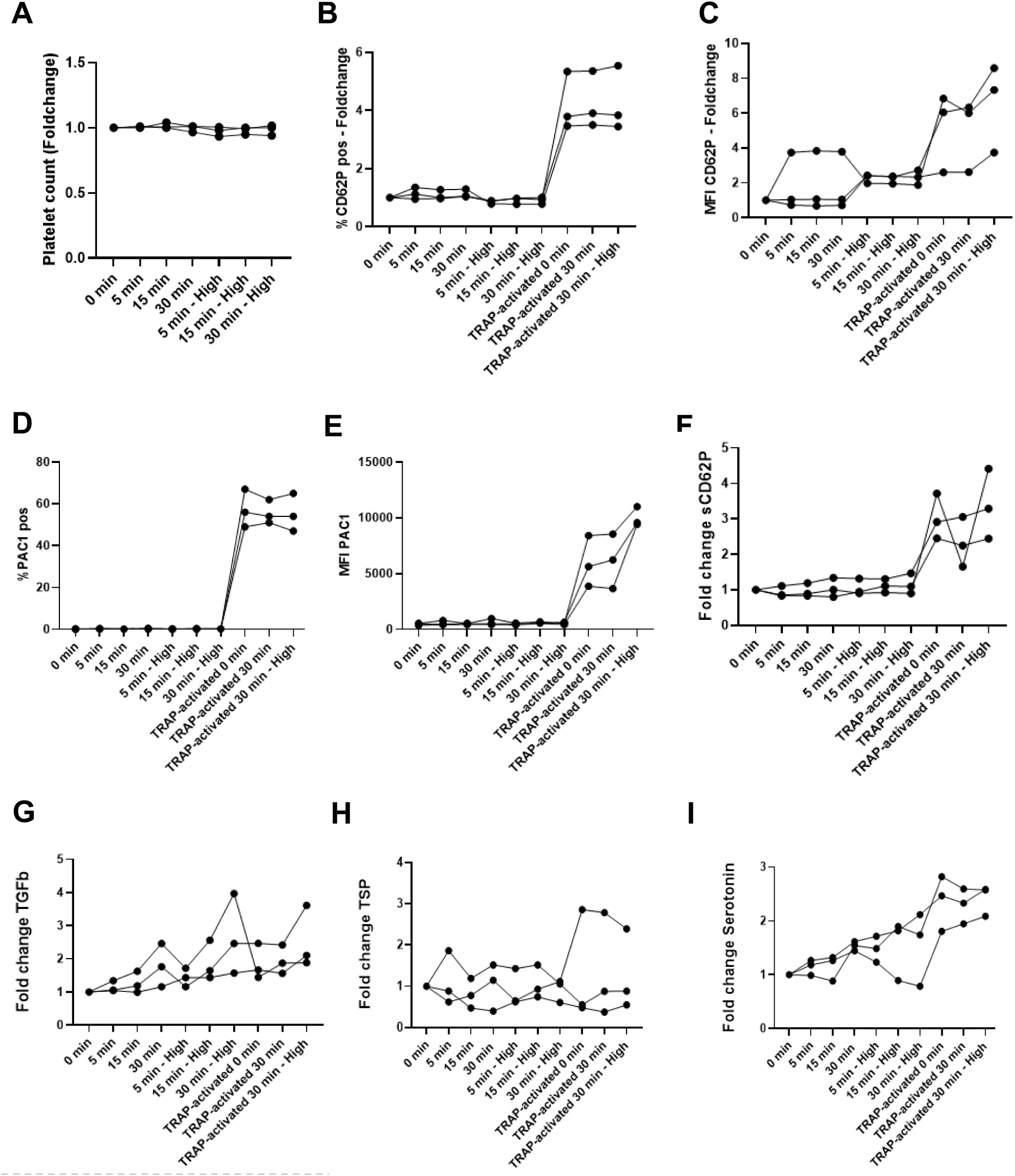
Analysis of platelet activation and degranulation after perfusion with apheresis platelet concentrate from a healthy donor. Samples were taken at 0, 5, 15, and 30 min under physiological (480 mL/min) and high (800 mL/min) flow. A) Platelet count (fold change). B) CD62P-positive platelets (% fold change). C) CD62P MFI (fold change). D) PAC1-positive platelets (%). E) PAC1 MFI. F) Foldchange of sCD62P. G) Foldchange of TGFβ. H) Foldchange of TSP. I) Foldchange of Serotonin.

Platelet membrane activation was assessed pre-/post-flow using CD62P and PAC-1 as a conformational marker of the GPIIb/IIIa integrin, which mediates platelet aggregation and platelet recruitment to ECs, and platelet irreversible activation. At both physiological and high flow rates, the percentage of CD62P-positive and PAC-1– positive platelets remained unchanged, indicating the absence of spontaneous activation and the maintenance of GPIIb/IIIa in its low affinity state. However, high flow was associated with a trend toward increased CD62P mean fluorescence intensity (MFI). TRAP-6 stimulation induced robust CD62P and PAC-1 responses, confirming platelet reactivity and validating the model for flow-induced activation studies. Together, these results indicate that neither physiological nor high flow rates triggered GPIIb/IIIa activation, while CD62P expression tended to rise under high flow, supporting the model’s suitability for platelet study (Figure 6B-E).

Degranulation was analyzed via ELISA of soluble factors (Figure 6F-I). Platelets under low and high flow showed progressive, shear-dependent increases in CD62P and TGF-β, consistent serotonin release, and variable TSP-1 release. These results suggest a local, flow-driven degranulation occurring without global platelet activation, highlighting secretion as a distinct platelet response to pathological hemodynamic conditions. These factors could exert local effects on ECs and SMC crosstalk, thereby contributing to stenosis development.

## Discussion

This study established a personalized real-scale CA model integrating patient-specific geometry, hemodynamic parameters and cellular behavior. The methodology is designed to bridge the gap between conventional *in vitro* systems and *in vivo* models, enabling more physiologically relevant investigations of vascular function and pathology.

The model consists of a personalized CA, fabricated via AM and adapted to defined biological constraints. It features a perfusable arterial channel lined with ECs as the cellular interface, integrated into a fluidic platform that allows controlled manipulation of hemodynamic parameters to reproduce physiological flow (the content). The system has been validated using existing imaging and diagnostic methods, and functional assays demonstrate endothelial responses and interactions with platelet components as a proof of concept.

Catano et al. combined AM, controlled hemodynamics, and ECs, showing for the first time the platform’s value for vascular pathologies. However, miniaturization scales flow but not ECs, exaggerating shear gradients and thus misrepresenting local cellular mechanics [44]. The model also lacks pulsatile flow, complex geometry, and integration with imaging or diagnostics.

Model accuracy has been discussed previously [26] and is mainly limited by MRI precision, which depends on patient factors, imaging parameters [45], and technical aspects of the MRI system [46]. Although arteriography is the gold standard, its use in SCD patients is limited.

The aM model is limited by its rigidity. Although compliant molds using PDMS or PVA-H exist [47], they do not match arterial compliance. In rigid models, absent damping exposes ECs to high pressure peaks, and the lack of cyclic stretch downregulates VE-cadherin, eNOS, and PECAM-1 [48].

The MCL uses a two-element Windkessel model (RC-circuit) to recreate pulsatile flow by adjusting flow rates and systolic/diastolic pressures, balancing experimental simplicity and physiological accuracy. Three-element models remain lumped parameter models, while four-element models add impedance and inertial elements to better capture pulsatile and inertial effects [29], [31], [30]. These adjustments could help correct the elevated diastolic values measured by Doppler but would greatly increase adjustment complexity, which must remain quick when cells are included.

HUVECs are chosen for their robust inflammatory responses, which are stronger than those of arterial ECs [35], though they differ structurally and functionally from arterial cells [32] with transcriptional heterogeneity [36]. Structural heterogeneity refers to variations in EC shape, size, and alignment, while functional heterogeneity reflects differences in their physiological roles and responses. Different EC types can be selected depending on the experimental purpose.

Gravitational seeding with manual rotation was used for its simplicity, though it is less efficient for 3D structures [49]. More effective methods include single-or dual-axis rotational seeding for tubular or complex geometries [38], [37], and magnetic seeding with cell-attached particles [40], [41].

ECs exhibited an early apoptotic phenotype, which preconditioning may help improve. Due to the model’s complex, non-tubular geometry, no shear preconditioning was applied, as variable flow directions could induce misalignment and promote pro-fibrotic or proliferative responses. Cells remained attached and aligned under flow, although shear may have initially stressed them. Applying shear to non-aligned ECs can transiently activate the NF-κB pathway and suppress eNOS activity until proper reorientation occurs [50]. Future studies could explore gradual pulsatile preconditioning to adapt ECs to shear stress and pressure [51]. However, EC alignment depends not only on shear stress but also on anisotropic strain [52], fiber orientation [53], and VEGF-A presence [54].

The decoupling of platelet degranulation and activation, previously observed under viral integration [55], suggests that localized release of profibrotic and proliferative factors such as TGF-β, TSP, and serotonin, in the absence of global activation, may influence EC–SMC crosstalk, promoting excessive SMC proliferation within the vascular intima and contributing to focal stenosis formation.

By enabling modular control over the container–content interface and accommodating multiple cell types, this platform offers a flexible, adaptable system that can be tailored to diverse pathologies and experimental conditions, providing a common foundation for translational research and personalized vascular modeling.

Future work will combine the model with Computational Fluid Dynamics (CFD) to map flow vectors and correlate them with EC alignment. Integration of hemolysis products and single-cell transcriptomics of ECs will enable analysis of inflammatory, oxidative, and mechanosensitive pathways. To investigate the interplay with SMCs, platelet and EC supernatants will be applied to assess proliferation, migration, and phenotypic transition. By systematically modulating pathological and healthy conditions, we aim to elucidate causal relationships between biological, geometric, and hemodynamic factors driving SCD-related cerebrovascular complications.

This work establishes a foundation for a new generation of personalized vascular models that bridge engineering precision and biological relevance. By integrating patient-specific geometry, pulsatile flow, and cellular components, the presented platform advances beyond conventional *in vitro* systems toward a controllable yet physiologically meaningful environment. Its adaptability enables mechanistic studies of all vascular pathologies which involve content-container interaction, such as SCD, atherosclerosis, and aneurysms, as well as the testing of medical devices under clinically representative conditions. Future developments focusing on material compliance and pathological flow patterns will further enhance its translational value. Ultimately, this approach offers a scalable pathway toward reducing animal experimentation, improving preclinical predictivity, and paving the way for individualized vascular therapies.

## References

[1] D. C. Rees, T. N. Williams, and M. T. Gladwin, “Sickle-cell disease,” The Lancet, vol. 376, no. 9757, pp. 2018–2031, Dec. 2010, doi: 10.1016/S0140-6736(10)61029-X.

[2] P. L. Kavanagh, T. A. Fasipe, and T. Wun, “Sickle Cell Disease: A Review,” JAMA, vol. 328, no. 1, p. 57, Jul. 2022, doi: 10.1001/jama.2022.10233.

[3] J. Z. Xu and S. L. Thein, “The carrier state for sickle cell disease is not completely harmless,” Haematologica, vol. 104, no. 6, pp. 1106–1111, Jun. 2019, doi: 10.3324/haematol.2018.206060.

[4] Ohene-Frempong, K et al., “Cerebrovascular Accidents in Sickle Cell Disease: Rates and Risk Factors,” *Blood*, The Journal of the American Society of Hematology, vol. Volume 91, Issue 1, pp. 288–294, Jan. 1998, doi: 10.1182/blood.V91.1.288.

[5] F. Bernaudin et al., “Chronic and acute anemia and extracranial internal carotid stenosis are risk factors for silent cerebral infarcts in sickle cell anemia,” Blood, vol. 125, no. 10, pp. 1653–1661, Mar. 2015, doi: 10.1182/blood-2014-09-599852.

[6] R. Adams et al., “The Use of Transcranial Ultrasonography to Predict Stroke in Sickle Cell Disease,” N Engl J Med, vol. 326, no. 9, pp. 605–610, Feb. 1992, doi: 10.1056/NEJM199202273260905.

[7] C. A. Hillery and J. A. Panepinto, “Pathophysiology of Stroke in Sickle Cell Disease,” Microcirculation, vol. 11, no. 2, pp. 195–208, Mar. 2004, doi: 10.1080/10739680490278600.

[8] P. Connes, S. Verlhac, and F. Bernaudin, “Advances in understanding the pathogenesis of cerebrovascular vasculopathy in sickle cell anaemia,” Br J Haematol, vol. 161, no. 4, pp. 484–498, May 2013, doi: 10.1111/bjh.12300.

[9] J. A. Switzer, D. C. Hess, F. T. Nichols, and R. J. Adams, “Pathophysiology and treatment of stroke in sickle-cell disease: present and future,” The Lancet Neurology, vol. 5, no. 6, pp. 501–512, Jun. 2006, doi: 10.1016/S1474-4422(06)70469-0.

[10] K. H. Merkel, P. L. Ginsberg, J. C. Parker, and M. J. Post, “Cerebrovascular disease in sickle cell anemia: a clinical, pathological and radiological correlation.,” Stroke, vol. 9, no. 1, pp. 45–52, Jan. 1978, doi: 10.1161/01.STR.9.1.45.

[11] K.-A. Nguyen-Peyre et al., “Impact of Carotid Geometry on Sickle Cell Diseases-Related Cerebral Vasculopthy Development,” Blood, vol. 140, no. Supplement 1, pp. 7820–7821, Nov. 2022, doi: 10.1182/blood-2022-156410.

[12] N. M. Oyesiku, D. L. Barrow, J. R. Eckman, S. C. Tindall, and A. R. T. Colohan, “Intracranial aneurysms in sickle-cell anemia: clinical features and pathogenesis,” Journal of Neurosurgery, vol. 75, no. 3, pp. 356–363, Sep. 1991, doi: 10.3171/jns.1991.75.3.0356.

[13] J. Singh, A. M. Ruhoff, D. Ashok, S. G. Wise, and A. Waterhouse, “Engineering advanced in vitro models of endothelial dysfunction,” Trends in Biotechnology, p. S0167779925000897, Apr. 2025, doi: 10.1016/j.tibtech.2025.03.004.

[14] S. Kamimura, M. Smith, S. Vogel, L. E. F. Almeida, S. L. Thein, and Z. M. N. Quezado, “Mouse models of sickle cell disease: Imperfect and yet very informative,” *Blood Cells*, Molecules, and Diseases, vol. 104, p. 102776, Jan. 2024, doi: 10.1016/j.bcmd.2023.102776.

[15] T. M. Franks et al., “Engineering, Generation and Preliminary Characterization of a Humanized Porcine Sickle Cell Disease Animal Model,” Sep. 16, 2020. doi: 10.1101/2020.09.15.291864.

[16] C. E. Kuczynski et al., “Novel Sheep Model of Sickle Cell Disease Reproduces Human Clinical and Laboratory Parameters,” Blood, vol. 140, no. Supplement 1, pp. 8220–8220, Nov. 2022, doi: 10.1182/blood-2022-168050.

[17] L. Goubergrits et al., “CT-based comparison of porcine, ovine, and human pulmonary arterial morphometry,” Sci Rep, vol. 13, no. 1, p. 20211, Nov. 2023, doi: 10.1038/s41598-023-47532-8.

[18] J. A. Catano et al., “A Patient-Specific 3D Printed Carotid Artery Model Integrating Vascular Structure, Flow, and Endothelium Responses,” Adv Healthcare Materials, p. e02478, Aug. 2025, doi: 10.1002/adhm.202502478.

[19] X. R. Lim and O. F. Harraz, “Mechanosensing by Vascular Endothelium,” Annu. Rev. Physiol., vol. 86, no. 1, pp. 71–97, Feb. 2024, doi: 10.1146/annurev-physiol-042022-030946.

[20] M. Nobili, J. Sheriff, U. Morbiducci, A. Redaelli, and D. Bluestein, “Platelet Activation Due to Hemodynamic Shear Stresses: Damage Accumulation Model and Comparison to In Vitro Measurements,” ASAIO Journal, vol. 54, no. 1, pp. 64–72, Jan. 2008, doi: 10.1097/MAT.0b013e31815d6898.

[21] T. Zhang et al., “The rapid change of shear rate gradient is beneficial to platelet activation,” Platelets, vol. 35, no. 1, p. 2288679, Dec. 2024, doi: 10.1080/09537104.2023.2288679.

[22] J. Sheriff, D. Bluestein, G. Girdhar, and J. Jesty, “High-Shear Stress Sensitizes Platelets to Subsequent Low-Shear Conditions,” Ann Biomed Eng, vol. 38, no. 4, pp. 1442–1450, Apr. 2010, doi: 10.1007/s10439-010-9936-2.

[23] P. Bartolucci, K.-A. Nguyen, C. Kassasseya, F. Segonds, M. Ritter, and P. Vedel, “Method for manufacturing a tridimensionnal blood vessel, WO2021/089665A1. W. I. P. Organization.”

[24] L. Václavů et al., “Intracranial 4D flow magnetic resonance imaging reveals altered haemodynamics in sickle cell disease,” Br J Haematol, vol. 180, no. 3, pp. 432–442, Feb. 2018, doi: 10.1111/bjh.15043.

[25] C. Renoux et al., “Effect of Age on Blood Rheology in Sickle Cell Anaemia and Sickle Cell Haemoglobin C Disease: A Cross-Sectional Study,” PLoS ONE, vol. 11, no. 6, p. e0158182, Jun. 2016, doi: 10.1371/journal.pone.0158182.

[26] S. Eckert et al., “Additive manufacturing of personalized scaffolds for vascular cell studies in large arteries: A case study on carotid arteries in sickle cell disease patients,”Annals of 3D Printed Medicine, vol. 16, p. 100178, Nov. 2024, doi: 10.1016/j.stlm.2024.100178.

[27] R. J. Adams et al., “Prevention of a First Stroke by Transfusions in Children with Sickle Cell Anemia and Abnormal Results on Transcranial Doppler Ultrasonography,” N Engl J Med, vol. 339, no. 1, pp. 5–11, Jul. 1998, doi: 10.1056/NEJM199807023390102.

[28] K. Sagawa, “Translation of Otto frank’s paper ‘Die Grundform des arteriellen Pulses’ zeitschrift für biologie 37: 483–526 (1899),” Journal of Molecular and Cellular Cardiology, vol. 22, no. 3, pp. 253–254, Mar. 1990, doi: 10.1016/0022-2828(90)91459-K.

[29] E. O. Kung and C. A. Taylor, “Development of a Physical Windkessel Module to Re-Create In Vivo Vascular Flow Impedance for In Vitro Experiments,” Cardiovasc Eng Tech, vol. 2, no. 1, pp. 2–14, Mar. 2011, doi: 10.1007/s13239-010-0030-6.

[30] N. Stergiopulos, J. J. Meister, and N. Westerhof, “Evaluation of methods for estimation of total arterial compliance,” American Journal of Physiology-Heart and Circulatory Physiology, vol. 268, no. 4, pp. H1540–H1548, Apr. 1995, doi: 10.1152/ajpheart.1995.268.4.H1540.

[31] N. Stergiopulos, B. E. Westerhof, and N. Westerhof, “Total arterial inertance as the fourth element of the windkessel model,” American Journal of Physiology-Heart and Circulatory Physiology, vol. 276, no. 1, pp. H81–H88, Jan. 1999, doi: 10.1152/ajpheart.1999.276.1.H81.

[32] W. C. Aird, “Phenotypic Heterogeneity of the Endothelium: I. Structure, Function, and Mechanisms,” Circulation Research, vol. 100, no. 2, pp. 158–173, Feb. 2007, doi: 10.1161/01.RES.0000255691.76142.4a.

[33] A. Przysinda, W. Feng, and G. Li, “Diversity of Organism-Wide and Organ-Specific Endothelial Cells,” Curr Cardiol Rep, vol. 22, no. 4, p. 19, Apr. 2020, doi: 10.1007/s11886-020-1275-9.

[34] N. D. Lawson and B. M. Weinstein, “Arteries and veins: making a difference with zebrafish,” Nat Rev Genet, vol. 3, no. 9, pp. 674–682, Sep. 2002, doi: 10.1038/nrg888.

[35] E. E. Eriksson, E. Karlof, K. Lundmark, P. Rotzius, U. Hedin, and X. Xie, “Powerful Inflammatory Properties of Large Vein Endothelium In Vivo,” ATVB, vol. 25, no. 4, pp. 723–728, Apr. 2005, doi: 10.1161/01.ATV.0000157578.51417.6f.

[36] J. Kalucka et al., “Single-Cell Transcriptome Atlas of Murine Endothelial Cells,” Cell, vol. 180, no. 4, pp. 764–779.e20, Feb. 2020, doi: 10.1016/j.cell.2020.01.015.

[37] H. R. Laube and M. Matthäus, “A new semi-automatic endothelial cell seeding technique for biological prosthetic heart valves,” Int J Artif Organs, vol. 24, no. 4, pp. 243–246, Apr. 2001, doi: 10.1177/039139880102400413.

[38] S. Hsu, I. Tsai, D. Lin, and D. C. Chen, “The effect of dynamic culture conditions on endothelial cell seeding and retention on small diameter polyurethane vascular grafts,” Medical Engineering & Physics, vol. 27, no. 3, pp. 267–272, Apr. 2005, doi: 10.1016/j.medengphy.2004.10.008.

[39] T. Dunkern et al., “A Novel Perfusion System for the Endothelialisation of PTFE Grafts Under Defined Flow,” European Journal of Vascular and Endovascular Surgery, vol. 18, no. 2, pp. 105–110, Aug. 1999, doi: 10.1053/ejvs.1999.0829.

[40] H. Perea, J. Aigner, U. Hopfner, and E. Wintermantel, “Direct Magnetic Tubular Cell Seeding: A Novel Approach for Vascular Tissue Engineering,” Cells Tissues Organs, vol. 183, no. 3, pp. 156–165, 2006, doi: 10.1159/000095989.

[41] H. Perea, J. Aigner, J. T. Heverhagen, U. Hopfner, and E. Wintermantel, “Vascular tissue engineering with magnetic nanoparticles: seeing deeper,” J Tissue Eng Regen Med, vol. 1, no. 4, pp. 318–321, Jul. 2007, doi: 10.1002/term.32.

[42] P. B. Van Wachem, J. W. S. Stronck, R. Koers-Zuideveld, F. Dijk, and C. R. H. Wildevuur, “Vacuum cell seeding: a new method for the fast application of an evenly distributed cell layer on porous vascular grafts,” Biomaterials, vol. 11, no. 8, pp. 602–606, Oct. 1990, doi: 10.1016/0142-9612(90)90086-6.

[43] L. Soletti, A. Nieponice, J. Guan, J. J. Stankus, W. R. Wagner, and D. A. Vorp, “A seeding device for tissue engineered tubular structures,” Biomaterials, vol. 27, no. 28, pp. 4863–4870, Oct. 2006, doi: 10.1016/j.biomaterials.2006.04.042.

[44] J. M. Dolan, H. Meng, S. Singh, R. Paluch, and J. Kolega, “High Fluid Shear Stress and Spatial Shear Stress Gradients Affect Endothelial Proliferation, Survival, and Alignment,” Ann Biomed Eng, vol. 39, no. 6, pp. 1620–1631, Jun. 2011, doi: 10.1007/s10439-011-0267-8.

[45] Jyoti, Dr. Bijendar Kumar Meena, Dr. Deepak Singh Phogat, Mohit Kumar Dahiya, “To Assess the Factors Effecting Image Quality in Magnetic Resonance Imaging at 1.5T,” IJRMST, vol. 17, no. 1, pp. 40–50, 2024, doi: 10.37648/ijrmst.v17i01.007.

[46] I. Robertson, “OPTIMAL MAGNETIC RESONANCE IMAGING OF THE BRAIN,” Vet Radiology Ultrasound, vol. 52, no. s1, Mar. 2011, doi: 10.1111/j.1740-8261.2010.01781.x.

[47] A. Shiravand et al., “Fabrication, characterization and numerical validation of a novel thin-wall hydrogel vessel model for cardiovascular research based on a patient-specific stenotic carotid artery bifurcation,” Sci Rep, vol. 14, no. 1, p. 16301, Jul. 2024, doi: 10.1038/s41598-024-66777-5.

[48] Y. Shi, D. Li, B. Yi, H. Tang, T. Xu, and Y. Zhang, “Physiological cyclic stretching potentiates the cell–cell junctions in vascular endothelial layer formed on aligned fiber substrate,” Biomaterials Advances, vol. 157, p. 213751, Feb. 2024, doi: 10.1016/j.bioadv.2023.213751.

[49] G. A. Villalona et al., “Cell-Seeding Techniques in Vascular Tissue Engineering,” Tissue Engineering Part B: Reviews, vol. 16, no. 3, pp. 341–350, Jun. 2010, doi: 10.1089/ten.teb.2009.0527.

[50] C. Wang, B. M. Baker, C. S. Chen, and M. A. Schwartz, “Endothelial Cell Sensing of Flow Direction,” ATVB, vol. 33, no. 9, pp. 2130–2136, Sep. 2013, doi: 10.1161/ATVBAHA.113.301826.

[51] R. Sodian et al., “Tissue-Engineering Bioreactors: A New Combined Cell-Seeding and Perfusion System for Vascular Tissue Engineering,” Tissue Engineering, vol. 8, no. 5, pp. 863–870, Oct. 2002, doi: 10.1089/10763270260424222.

[52] S. Iwayoshi, K. Furukawa, and T. Ushida, “Continuous Visualization of Morphological Changes in Endothelial Cells in Response to Cyclic Stretch,” *JSME Int. J.*, Ser. C, vol. 49, no. 2, pp. 545–555, 2006, doi: 10.1299/jsmec.49.545.

[53] Y. Li et al., “Engineering cell alignment in vitro,” Biotechnology Advances, vol. 32, no. 2, pp. 347–365, Mar. 2014, doi: 10.1016/j.biotechadv.2013.11.007.

[54] A.-C. Vion et al., “Endothelial Cell Orientation and Polarity Are Controlled by Shear Stress and VEGF Through Distinct Signaling Pathways,” Front. Physiol., vol. 11, p. 623769, Mar. 2021, doi: 10.3389/fphys.2020.623769.

[55] L. J. Weiss et al., “Uncoupling of platelet granule release and integrin activation suggests GPIIb/IIIa as a therapeutic target in COVID-19,” Blood Advances, vol. 7, no. 11, pp. 2324–2338, Jun. 2023, doi: 10.1182/bloodadvances.2022008666.

